# Neonatal brain-directed gene therapy rescues a mouse model of neurodegenerative CLN6 Batten disease

**DOI:** 10.1101/673848

**Authors:** Sophia-Martha kleine Holthaus, Saul Martin-Herranz, Giulia Massaro, Mikel Aristorena, Justin Hoke, Michael P. Hughes, Ryea Maswood, Olha Semenyuk, Mark Basche, Amna Z. Shah, Izabela P. Klaska, Alexander J. Smith, Sara E. Mole, Ahad A Rahim, Robin R Ali

## Abstract

The neuronal ceroid lipofuscinoses (NCLs), more commonly referred to as Batten disease, are a group of inherited lysosomal storage disorders that present with neurodegeneration, loss of vision and premature death. There are at least 13 genetically distinct forms of NCL. Enzyme replacement therapies and preclinical studies on gene supplementation have shown promising results for NCLs caused by lysosomal enzyme deficiencies. The development of gene therapies targeting the brain for NCLs caused by defects in transmembrane proteins has been more challenging and only limited therapeutic effects in animal models have been achieved so far. Here, we describe the development of an adeno-associated virus (AAV)-mediated gene therapy to treat the neurodegeneration in a mouse model of CLN6 disease, a form of NCL with a deficiency in the membrane-bound protein CLN6. We show that neonatal bilateral intracerebroventricular injections with AAV9 carrying *CLN6* increase lifespan by more than 90%, maintain motor skills and motor coordination and reduce neuropathological hallmarks of *Cln6*-deficient mice up to 23 months post vector administration. These data demonstrate that brain-directed gene therapy is a valid strategy to treat the neurodegeneration of CLN6 disease and may be applied to other forms of NCL caused by transmembrane protein deficiencies in the future.

## Introduction

The neuronal ceroid lipofuscinoses (NCLs), also known as Batten disease, are a group of fatal, inherited neurodegenerative lysosomal storage diseases (LSDs). They are characterised by the progressive accumulation of autofluorescent storage material in the lysosome. Clinical symptoms of NCL usually present in childhood and include loss of vision, seizures, cognitive decline and mental deterioration ultimately resulting in premature death (1). The combined incidence of NCL varies across different countries from 1:12,500 to 1:100,000(2). At least 13 genes are linked to the development of the disease, five of which encode transmembrane proteins that reside in the membrane of the lysosome, endoplasmic reticulum (ER) or ER-Golgi intermediate compartment(3). There is no licensed therapy available for NCLs caused by transmembrane protein deficiencies and only palliative care is offered to patients.

CLN6 disease is a form of NCL caused by mutations in the gene *CLN6* encoding a transmembrane, endoplasmic reticulum (ER) protein of unknown function(4). Disease symptoms usually present in children between 1.5 to 8 years of age and include loss of vision, ataxia, motor delay, seizures and cognitive abnormalities. Childhood CLN6 disease progresses rapidly leading to premature death between 5 to 12 years of age (5). Mutations in *CLN6* can also lead to a rare adult-onset form of NCL, referred to as Kufs disease. The main differences between the two conditions are the latter age of onset and a lack of visual involvement in adult patients(6).

The *Cln6*^*nclf*^ mouse is a naturally occurring model of CLN6 disease. *Cln6*^*nclf*^ mice carry a 1-bp insertion mutation in the gene *Cln6*. It is analogous to a common mutation found in CLN6 patients of Pakistani origin that results in a premature stop codon and a truncated protein product(7, 8). *Cln6*-deficient mice recapitulate the human disease with severe retinal and neuronal degeneration. The disease phenotype involves blindness, progressive loss of locomotor function and motor coordination, and a reduced mean survival of approximately 12 months of age, rendering it a useful animal model to study therapeutic interventions for CLN6 disease(9).

Adeno-associated virus (AAV)-mediated gene therapies are proving to be effective for the treatment of a wide range of inherited disorders(10–14). However, the development of gene therapies for conditions that affect multiple regions of the brain appears to be more challenging due to its large size, complex anatomy and the presence of the blood brain barrier (BBB). For this reason, AAV-mediated gene therapy clinical trials for neurodegenerative LSDs have mostly focused on conditions involving genes encoding soluble enzymes including a safety phase I clinical trial for late-infantile onset CLN2 disease (NCT00151216)(15). Therapies for these diseases are aided by lysosomal cross-correction allowing therapeutic enzymes to be secreted and passed on to surrounding cells via the mannose-6-phosphate pathway(16). To achieve meaningful therapeutic effects for transmembrane protein deficiencies is more challenging as cross-correction does not occur, confining benefit to only those cells that received the vector(17, 18).

A phase I/II clinical trial involving intrathecal delivery of a self-complementary (sc) AAV9.Cb.CLN6 to children suffering from CLN6 disease has been initiated recently (NCT02725580). However, no pre-clinical or clinical data has been published to date demonstrating that CNS-directed gene therapy is beneficial in CLN6 disease. Previously, we have demonstrated that AAV gene supplementation therapy can be used to treat the ocular phenotype in *Cln6*^*nclf*^ mice(19). Here, we report the successful development of an AAV-mediated gene therapy directly targeting the brain of new-born, presymptomatic *Cln6*^*nclf*^ mice. We establish that neonatal bilateral intracerebroventricular (ICV) injections with an AAV9 vector carrying human *CLN6* extend lifespan by more than 90%, attenuate motor skill and coordination deficits, and reduce the neuropathological hallmarks evident in mutant mice. Our work demonstrates the long-term therapeutic effects of AAV-mediated gene therapy in a model of CLN6 disease, and supports the development of CNS-directed gene therapies for CLN6 disease and other forms of NCL caused by transmembrane protein deficiencies.

## Results

### Bilateral ICV delivery of AAV9.CMV.hCLN6 leads to a widespread expression of *CLN6* throughout the brains of wild type mice

Previously, mRNA expression analyses showed *Cln6* expression in wild type mouse brains across various regions including cerebellum, hypothalamus, mid brain, cortex (frontal, posterior and entorhinal), hippocampus, thalamus and striatum. Expression was detected from embryonic day (E) 18 with significantly lower Cln6 mRNA levels in *Cln6*^*nclf*^ mutant brains than in wild type brains(20). Based on these data, we performed bilateral ICV injections in wild type P0-P1 mice (under 24h of age) with a single stranded AAV9 vector carrying human CLN6 (hCLN6) under the control of the CMV promoter (AAV9.CMV.hCLN6). A low dose of 5×10^10^ viral genomes (vg) (n = 3) or a high dose of 5×10^11^ vg (n = 3) were administered per pup. Non-injected littermates were used as age-matched controls (n = 3). Four weeks post administration, the mouse brains were harvested, fixed and cryo-sectioned along the coronal axis to investigate the biodistribution and transduction efficiency. Immunostaining for human CLN6 4 revealed an extensive rostrocaudal *CLN6* expression throughout the brain following injections with the high dose of the vector. Representative images are shown from the cortex (somatosensory cortex barrel field), striatum (caudate nucleus), hippocampus (CA2-3), thalamus (ventral posterolateral and postermedial nuclei), cerebellum (central lobe) and brainstem (gigantocellular nucleus). While administration of the low dose also led to widespread *CLN6* expression, anti-CLN6 staining was weaker and regions of the brains were only sparsely transduced (e.g. cortex, cerebellum) (Figure 1). Higher magnification light microscopy of discrete regions of the brain sections revealed CLN6 staining in cells with both neuronal and glial morphology.

**Fig. 1.**
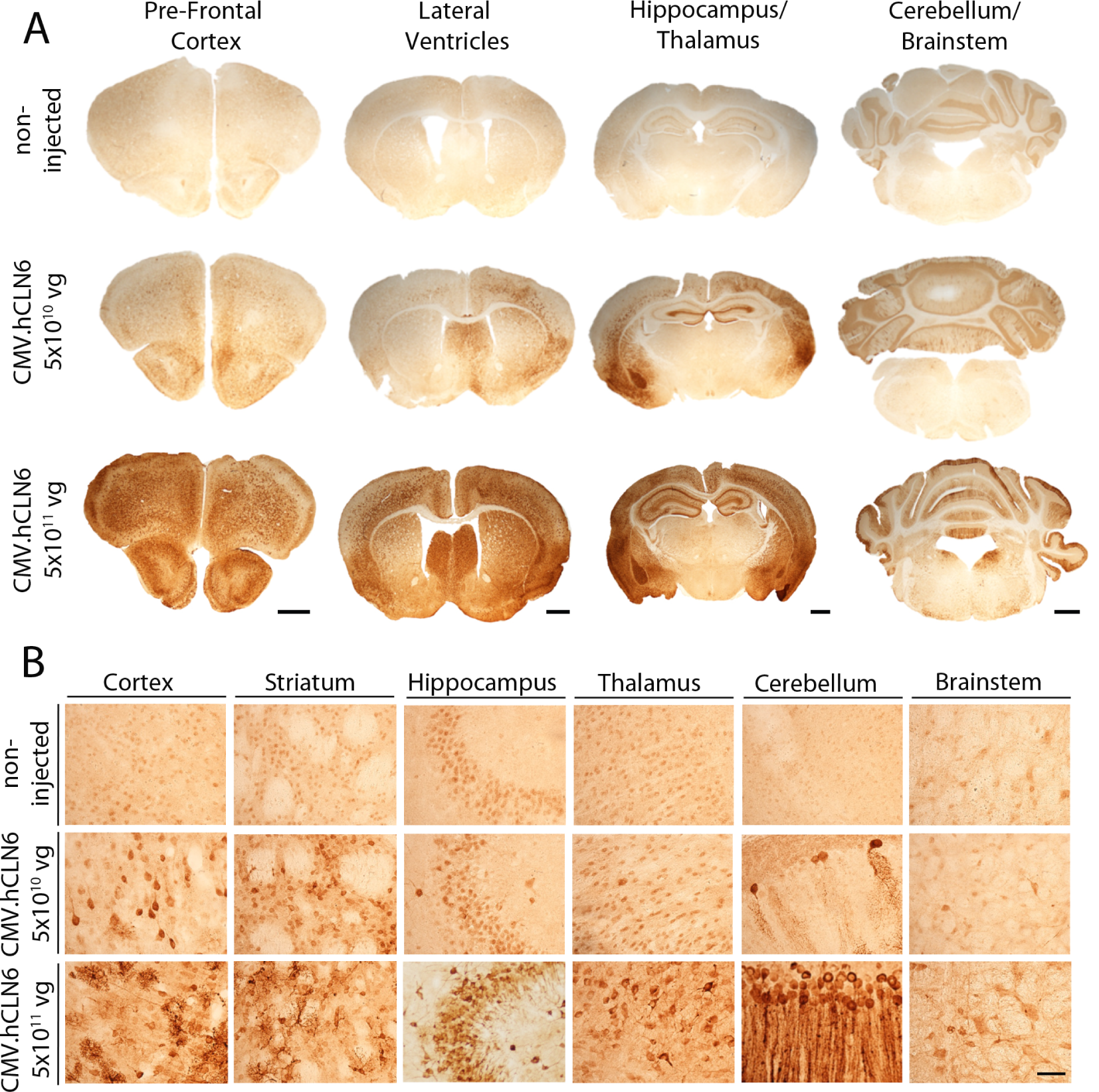
Neonatal ICV injections of AAV9.CMV.hCLN6 lead to widespread expression of *CLN6* in wild type mice. (A) Four weeks post-administration, widespread rostrocaudal expression of *CLN6* was detected throughout the brains of wild type mice that received AAV9.CMV.hCLN6 at 5×10^11^ vg (n = 3). Lower transduction efficiency and expression levels were observed in wild type mice that received vector at 5×10^10^ vg (n = 3). CLN6 staining was not detected in untreated wild type controls (n = 3). (B) Higher magnification images of CLN6-positive cells across various brain regions following low and high titre administration of AAV9.CMV.hCLN6. Scale bars (A): 1mm, (B): 100 µm.

To assess for any astroglial or microglial-mediated inflammatory responses associated with overexpression of human CLN6 in wild type mice, immunostaining with antibodies against the glial fibrillary acidic protein (GFAP) and CD68 was performed. An examination of brain sections stained for GFAP revealed no differences in astrocyte cell morphology and no increase in GFAP staining was observed in brains that received low dose vector compared with age-matched uninjected controls (Figure S1A, B). However, a subtle, confined but not significant up-regulation of GFAP staining was observed in the somatosensory barrel cortex region (S1BF) (Figure S1B, S3A). No increase in GFAP staining was detected in any other discrete region examined (Figure S1A, B). CD68 immunostaining did not reveal any obvious microglia-mediated inflammatory responses in wild type brains treated with the high or low dose compared with uninjected brains (Figure S2A-B, S3B).

### Brain-directed gene therapy normalises lifespan and preserves brain weight in *Cln6*^*nclf*^ *mice*

New-born (P0-P1) *Cln6*^*nclf*^ mice received bilateral ICV injections with AAV9.CMV.hCLN6 at 5×10^11^ vg (n = 6) and a small cohort of mutant mice received injections with a lower dose of 1×10^11^ vg (n = 2). Age-matched uninjected mutant mice (n = 14) and wild type mice were used as controls (n = 8). As mutant mice do not show a significant loss of bodyweight compared with age-matched wild type mice (Figure S3), the humane endpoint for the experiment was defined by either a poor score in the neurological welfare scoring system or a loss of 10-15% in bodyweight. Untreated *Cln6*^*nclf*^ mice had a mean survival of 11.7 ± 0.15 months with a maximum survival of 12 months (Figure 2A). Mutant mice that received AAV9.CMV.hCLN6 at 5×0^11^ vg and 1×10^11^ vg had a significantly prolonged survival with a mean of 22.7 ± 0.81 months (p = 0.0002) and 23.3 months (p = 0.0114), respectively. Three mice treated with high dose vector, one mouse treated with low dose vector and five wild type mice did not deteriorate to reach the humane endpoint. These mice were euthanised between 23 and 24 months of age (Figure 2A, S4C). An example video of an untreated end-stage mutant and age-matched treated mutant mouse is shown in supplementary video S1.

**Fig. 2.**
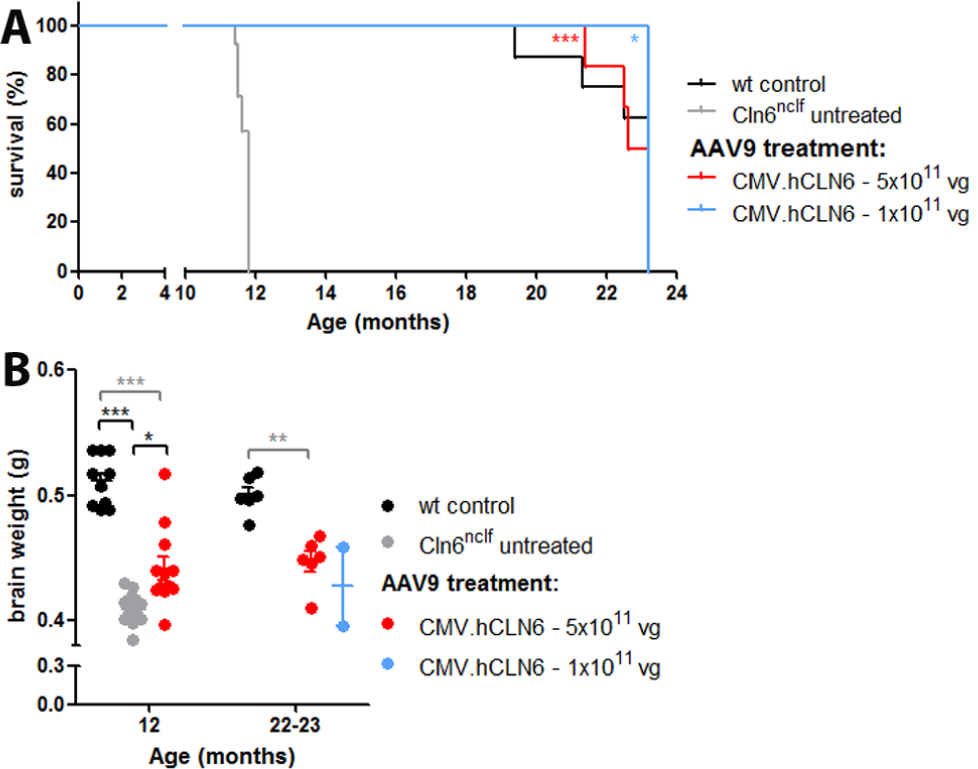
Neonatal ICV injections of AAV9.CMV.hCLN6 significantly prolongs survival and preserves brain weight of *Cln6*^*nclf*^ mice. (A) Kaplan-Meier survival curve showing percentage of survival over time. Median survival of mice was as follows: *Cln6*^*nclf*^ untreated, 11.7 ± 0.15 months (n = 14); wild type controls (n = 8); *Cln6*^*nclf*^ treated with 5×10^11^ vg AAV9.CMV.hCLN6, 22.7 ± 0.81 months (n = 6); *Cln6*^*nclf*^ treated 1×10^11^ vg AAV9.CMV.hCLN6, 23.3 months (n = 2). Data represented as means ± SEM, compared by Log-rank test (Mantel-Cox) (***p = 0.0002, *p = 0.0114). (B) Mean brain weights: wild type controls at 12 months, 0.511 ± 0.02 g (n = 10) and 22-23 months, 0.5 ± 0.01 g (n = 6); *Cln6*^*nclf*^ untreated, 0.410 ± 0.01 g (n = 20); *Cln6*^*nclf*^ treated with 5×10^11^ vg AAV9.CMV.hCLN6 at 12 months, 0.4413 ± 0.03g (n = 12) and 22-23 months, 0.454 ± 0.01 g (n = 6); 22-23 months *Cln6*^*nclf*^ treated with 1×10^11^ vg AAV9.CMV.hCLN6, 0.427 ± 0.04 g (n = 2). Data represented as means ± SD, compared by one-way ANOVA (black asterisks) and two-way ANOVA (grey asterisks) and Dunn’s multiple comparison test (* p < 0.05, *** p < 0.0001).

We measured the whole brain weight of age-matched treated, untreated and wild type mice. At 12 months, untreated mutant mice had significantly lighter brains than wild type animals (0.410 ± 0.01 g vs. 0.511 ± 0.02 g; p < 0.0001). Mutant mice that received bilateral ICV injections with AAV9.CMV.hCLN6 at 5×10^11^ vg had higher brain weights at 12 months than untreated age-matched mutants (0.4413 ± 0.03 g; p < 0.05). The brain weight of treated mice examined at 22-23 months was not significantly different compared with 12 months old treated brains (0.454 ± 0.01 g; p < 0.05). The weight of the treated brains was significantly lower than wild type brains at both 12 and 22-23 months (Figure 2B).

### ICV treatment improves motor skill and coordination deficits in *Cln6*^*nclf*^ mice

From 7 months of age, *Cln6*^*nclf*^ mice showed a progressive reduction in the latency to fall in an accelerating rotarod test compared with wild type mice (p < 0.01). Following treatment with high dose vector, mutant mice had a significantly better rotarod performance at 7 months compared with age-matched untreated mutant mice (p < 0.01). The performance of treated mutant mice was variable thereafter but remained better up to 12 months (Figure 3A). Motor skills and coordination were also assessed in a 1-minute foot-fault test. The number of times a limb fell through the grid was counted and the percentage of incorrect steps was calculated. Untreated *Cln6*-deficient mice made significantly more mistakes (∼9% incorrect steps at 9-11 months, ∼12% incorrect steps at 11.5 months) compared with wild type mice (∼6-8% incorrect steps from 9-months). Treated *Cln6*^*nclf*^ mice made significantly fewer mistakes than untreated mice (∼6-7% incorrect steps across all time points) and were comparable with age-matched wild type controls, demonstrating that the treatment normalised the foot-fault phenotype (Figure 3B).

**Fig. 3.**
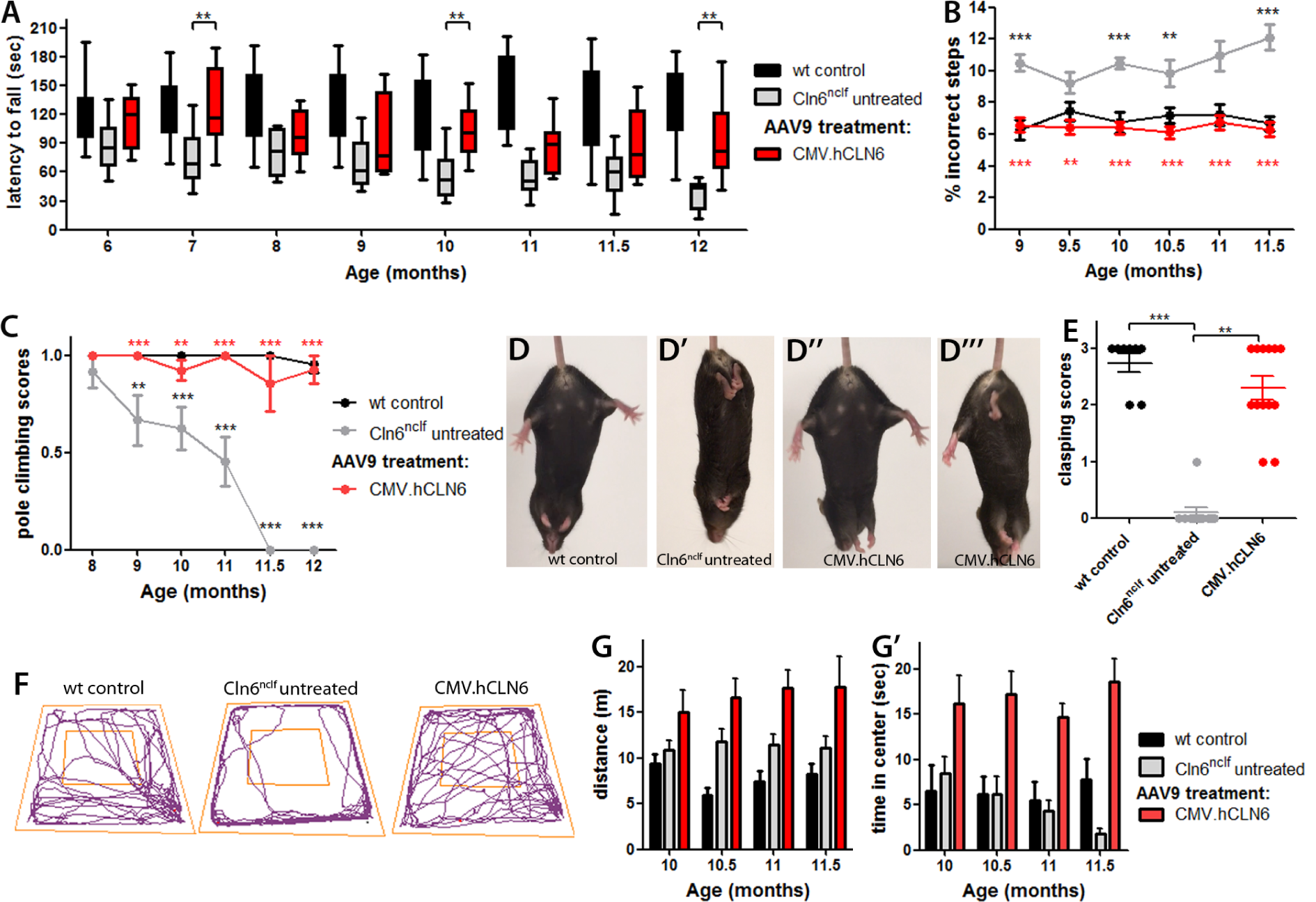
Neonatal ICV administration of AAV9.CMV.hCLN6 attenuates behavioural phenotypes of *Cln6*-deficient mice. (A) Accelerating rotarod performance measured as latency to fall, (B) 1-minute grid test performance measured as percentage of incorrect steps and (C) vertical pole climbing scores of untreated mutants, treated mutants and wild type controls over time. (D-D”‘) Example images of hind limp clasping behaviour and (E) quantification of hind limp clasping scores with untreated mutants (11.5-12 months), treated mutants (12 months) and wild type controls (12 months). Wild type controls, n = 10; *Cln6*^*nclf*^ untreated, n = 10 (7-11.5 months), n = 7, (12 months); *Cln6*^*nclf*^ treated, n = 13. Data are presented as (A) median (line), 25-75 percentile (box) and 10-90 percentile (whiskers) and as (B-E) means ± SEM, compared by two-way ANOVA and Bonferroni test with untreated *Cln6*^*nclf*^ as comparison group (*p < 0.05, **p < 0.01, ***p < 0.001; (B-C) black asterisks: wt vs. untreated, red asterisks: treated vs. untreated). (F) Representative open field test behaviour of different groups, (G) analysis of distance travelled and (G’) time spent in the centre of the arena in open field test over time. Wild type controls, n = 9-11;*Cln6*^*nclf*^ untreated, n = 11; *Cln6*^*nclf*^ treated, n = 14. Data are represented as means ± SEM and was compared by linear regression analysis.

To further assess motor function and co-ordination we used a vertical pole test. At 8 months, untreated mutants and wild type mice descended the pole without difficulties. From 9 months onwards, mutants struggled to perform the task and at 11.5 and 12 months they were unable to grasp and climb down the pole in a coordinated manner (see supplementary videos S2-S4). Treated mice performed significantly better than untreated animals and no significant differences were detected between treated *Cln6*^*nclf*^ mice and age-matched wild type controls (Figure 3C).

A 30-second tail suspension test was used to assess the hindleg clasping phenotype of the mice. In line with the pole climbing data, *Cln6*-deficient mice showed consistently severe hind limb clasping at 11.5 and 12 months of age (Figure 3D’,3E), which was not present in wild type mice (Figure 3D). Scoring of the phenotype by observers masked to each experimental cohort confirmed that clasping was significantly more severe in untreated *Cln6*^*nclf*^ mice (Figure 3E, p < 0.0001). At this age, treated *Cln6*^*nclf*^ mice had either no clasping phenotype (Figure 3D”) or a mild clasping phenotype (Figure 3D”‘) demonstrating that the treatment was effective (Figure 3E, p < 0.005).

Finally, we tested the mobility of the mice in an open field activity test (Figure 3F). Consistent with a previous report(9), no significant differences were found in the total distance travelled (Figure 3G), speed (Figure S5A, B), time immobile (Figure S5C) or time mobile (Figure S5D) of wild type and untreated *Cln6*^*nclf*^ mice over time. However, compared with wild type mice, untreated mutant mice visited the centre of the arena less frequently (for examples see Figure 3F) and spent less time in the centre than wild type mice (Figure 3G’). Linear regression analysis showed that the amount of time spent in the centre of the arena by the untreated animals decreased significantly between 10 and 11.5 months (p = 0.0059), while no significant change over time was detected in the distance travelled or the time spent in the centre by the treated *Cln6*^*nclf*^ mice or the wild type mice (Figure 3G, 3G’). Interestingly, this experiment did reveal that treated *Cln6*^*nclf*^ mice were overall more active than untreated and wild type mice, as shown by the longer distance travelled (Figure 3G) and more time spent in the centre (Figure 3G’).

### AAV9.CMV.hCLN6 treatment leads to long-term therapeutic benefit

To assess the long-term effects of the treatment, a cohort of mutant mice that received 5 × 10^11^ vg vector (n = 6) were kept beyond 12 months of age and the performance in the 1-minute foot-fault test was tested every month until the end of the study. The quantification of the test showed that treated *Cln6*^*nclf*^ mice performed the task as well as wild type mice up to 23 months of age (Figure 4A), although there may be a slight trend towards deterioration of mobility at the later time points. The performance of the mice was also assessed in the tail suspension test. No significant difference in hind limb clasping was detected between 22-23 months old treated mice and age-matched wild type controls (p = 0.417) (Figure 4B). In comparison with the performance at 12 months (Figure 3E), the average score of treated *Cln6*^*nclf*^ mice and wild type mice decreased by a similar extent over time, indicating that the treatment effect was maintained for almost 2 years following the injections, equivalent to the lifespan of wild type mice. It should be noted however, that the number of animals at the last time points is limited by the natural lifespan of both groups of mice. As a result the study is powered to detect only a greater than 50% difference in mean clasping score based on the current variation.

**Fig. 4.**
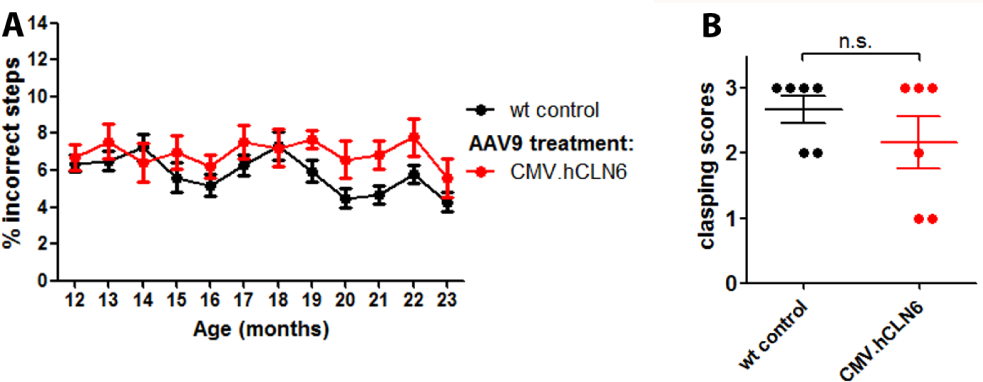
Treated *Cln6*^*nclf*^ mice show long-term correction of behavioural phenotypes. The performance of treated *Cln6*^*nclf*^ and wild type mice was (A) measured in the 1-minute foot-fault test over time and (B) scored in the hind limb clasping test at 22-23 months of age. No significant differences were detected in the two tests between treated mice and wild type controls. (A) Wild type, n = 8 (12-19 months), n = 6 (19-22 months), n = 5 (23 months); *Cln6*^*nclf*^ treated mice, n = 6 (12-21 months), n = 5 (23 months). (B) Wild type controls n = 6, *Cln6*^*nclf*^ treated mice n = 6. Data sets are represented as means ± SEM, compared by (A) two-way ANOVA and (B) non-parametric Mann-Whitney test.

### Brain-directed gene therapy preserves cortical thickness and reduces loss of neurons in *Cln6*-deficient mice

The brains of the mice were harvested, cut along the longitudinal fissure and the right hemisphere of each brain was fixed and cryo-sectioned along the coronal axis. To confirm the transgene expression in the brain of treated mice, we stained the brain sections with antibodies against human CLN6 and the neuronal nuclear marker NeuN. We observed sustained widespread expression of CLN6 throughout the cortex, thalamus, hippocampus (CA1 region) and the primary and secondary visual cortex (V1/V2 region) co-localising with a large number of NeuN-positive cells in treated brains at 12 and 22-23 months (Figure S6A, B). Co-staining with antibodies against GFAP and CD68 identified a small number of astrocytes that were also positive for CLN6 in the cortex, whereas we did not observe *CLN6* expression in microglia (Figure S6C). To quantify the number of transduced neurons, astrocytes and microglia, we counted the number of NeuN-, GFAP- and CD68-positive cells that were also positive for CLN6 staining in cortical region using the software image J. We found that in 12 months old brain sections approximately 50 percent of NeuN-, 20 percent of GFAP- and less than 5 percent of CD68-positive cells expressed *CLN6* (Figure S7). We also analysed the mRNA level of CLN6 across different brain regions in wild type, untreated mutant and treated mutant mice at 12 months of age. A direct comparison of the absolute number of mRNA molecules by real-time qPCR showed that the CLN6 expression levels were significantly higher in the cortex of treated mutants than the endogenous Cln6 levels in the cortex of wild type mice. No significant difference was detected between wild type and treated mutants in the visual cortex, hippocampus and thalamus (Figure S8). Neuronal loss and brain atrophy is a major hallmark of CLN6 disease that is recapitulated in *Cln6*-deficient mice with a reduced total brain weight and loss in cortical thickness (9, 21). To assess the effect of gene therapy on the neurodegeneration in *Cln6*^*nclf*^ mice, brain sections were immunostained with an antibody against NeuN to measure cortical thickness and to enable neuronal cell counts. The S1BF region was slightly thinner in untreated *Cln6*^*nclf*^ mouse brains at 12 months compared with age-matched wild type controls (1889 ± 8.84 µm vs. 2075 ± 59.14 µm; p = 0.01). We did not observe a rescue of cortical thickness following treatment (1968 ± 19.10 µm). Of note, no significant further loss in thickness was detected between 12 and 22-23 months of age (1918 ± 29.81 µm) (Figure 5A-B’). We assessed the number of neurons by stereological counts of NeuN-positive cells in the S1BF cortical region and the VPM/VPL nuclei of the thalamus, a brain region that presents with early disease pathology in *Cln6*^*nclf*^ mice (9, 21).In comparison with 12 month old wild type controls, untreated end-stage mice showed a significant loss of neurons in the S1BF region by 28% (118,980 ± 8,116 vs. 85,680 ± 2,839; p = 0.0064) and in the VPM/VPL region by 40% (37,020 ± 3,187 vs. 22,380 ± 2,613; p = 0.011). Neuronal cell counts of treated *Cln6*^*nclf*^ mouse brains at 12 months revealed significantly higher numbers of neurons compared with untreated brains, with a 23% increase in the S1BF region (113,700 ± 8,177; p = 0.0296) and a 30% increase in the VPM/VPL region (33,300 ± 2,076; p = 0.048). The analysis of treated *Cln6*^*nclf*^ mouse brains at 22-23 months showed no significant loss of neurons in the S1BF region (101,115 ± 4422; p = 0.4691) and VPM/VPL regions (29,280 ± 2196; p = 0.654) compared with 12 month-old treated mice (Figure 5B, B”, C-C’). These data demonstrate that neonatal ICV injection of AAV9.CMV.hCLN6 provide a significant and longterm amelioration of the neurodegeneration in *Cln6*^*nclf*^ mice.

**Fig. 5.**
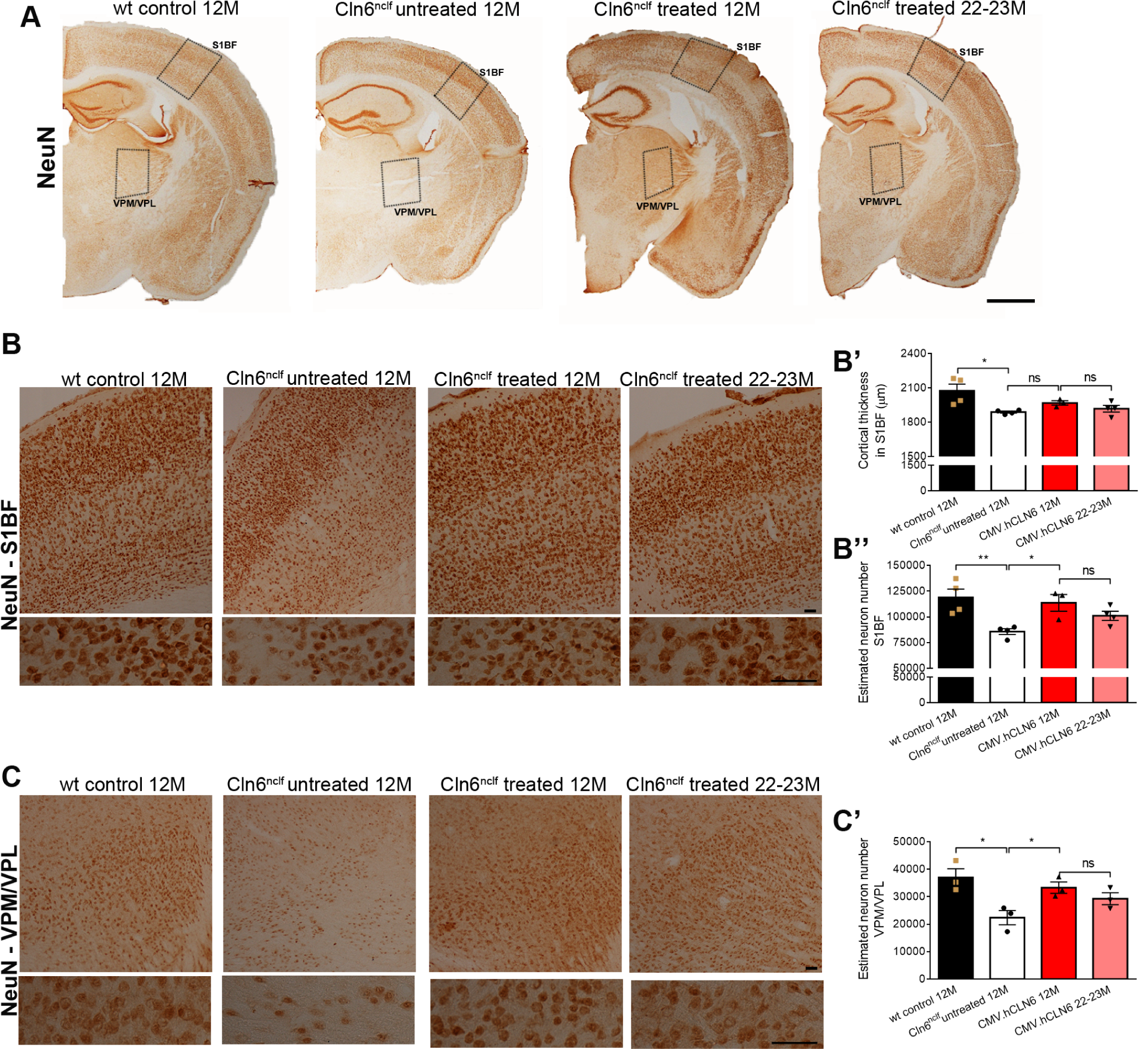
Brain-directed gene therapy preserves neuronal cells in *Cln6*^*nclf*^ mice long term. (A) Representative images of coronal brain cross-sections from 12-month-old end-stage Cln6^nclf^ mice, 12-month-old wild type mice, and 12- and 22-23-month-old treated *Cln6*^*nclf*^ mice. (B, C) Higher magnification images from S1BF and VPM/VPL regions stained for NeuN. (B’) Measurement of the cortical thickness in the S1BF region (B”, C’) and neuronal counts in the S1BF and VPM/VPL regions showing a significantly higher number of neurons in brains of *Cln6*^*nclf*^ mice at 12 months following neonatal ICV injections of vector. No significant difference was detected in the number of neurons in the S1BF and VPM/VPL region between brain sections of treated mutant mice aged 12 and 22-23 months. Scale bars: (A) 1 mm, (B, C) 50 µm. Wild type controls (12 months), n = 4; untreated *Cln6*^*nclf*^ (12 months), n = 4; treated *Cln6*^*nclf*^ (12 months), n = 3; treated *Cln6*^*nclf*^ mice (22-23 months), n = 4. Data are presented as means ± SEM and analysed by one-way ANOVA and Sidak’s multiple comparisons test (*p < 0.05, **p < 0.01).

### AAV9.CMV.hCLN6 treatment prevents lysosomal pathology and the accumulation of autofluorescent storage material in mutant mice

The progressive accumulation of autofluorescent storage material in the lysosome is a major neuropathological hallmark of CLN6 disease and prominent lysosomal inclusions were reported across several brain regions in *Cln6*-deficient mice including S1BF, VPM/VPL and V1/V2 region (9, 21). To assess lysosomal pathology and disease progression, we examined autofluorescence in the 488nm excitation channel and performed immunostaining against lysosomal-associated membrane protein 1 (LAMP1), a marker for late-endosomes and lysosomes. We found increased autofluorescence was consistently present close to the nucleus (Figure 6A) and increased LAMP1 immunostaining with larger LAMP1-positive areas in the S1BF, VPM/VPL and V1/V2 regions in end-stage mutant brain sections compared with age-matched wild type controls (Figure 6B). The autofluorescence was reduced and LAMP1-positive accumulations appeared to be fewer and smaller in brain sections from treated mice at 12 months and 22-23 months (Figure 6). To quantify LAMP1 immunostaining, thresholding immunoreactivity analysis was performed, excluding LAMP1-positive areas below 0.01 µm^2^ to focus on abnormally enlarged lysosomes. Untreated end-stage mutant brain sections had significantly larger areas of LAMP1-positive immunofluorescence across all brain regions investigated compared with age-matched wild type controls (p < 0.001 for S1BF, VPM/VPL and V1/V2). Treated *Cln6*^*nclf*^ mice had significantly smaller areas of LAMP1-positive immunofluorescence across all regions at 12 months (p < 0.001 for S1BF, VPM/VPL and V1/V2). No significant differences were detected between sections from treated mice at 12 and 22-23 months (Figure 6B’-B”’), highlighting the long-term therapeutic effect of the treatment. These data suggest that treatment reduces the disease-related lysosomal pathology and the accumulation of intracellular storage material in the brains of *Cln6*^*nclf*^ mice.

**Fig. 6.**
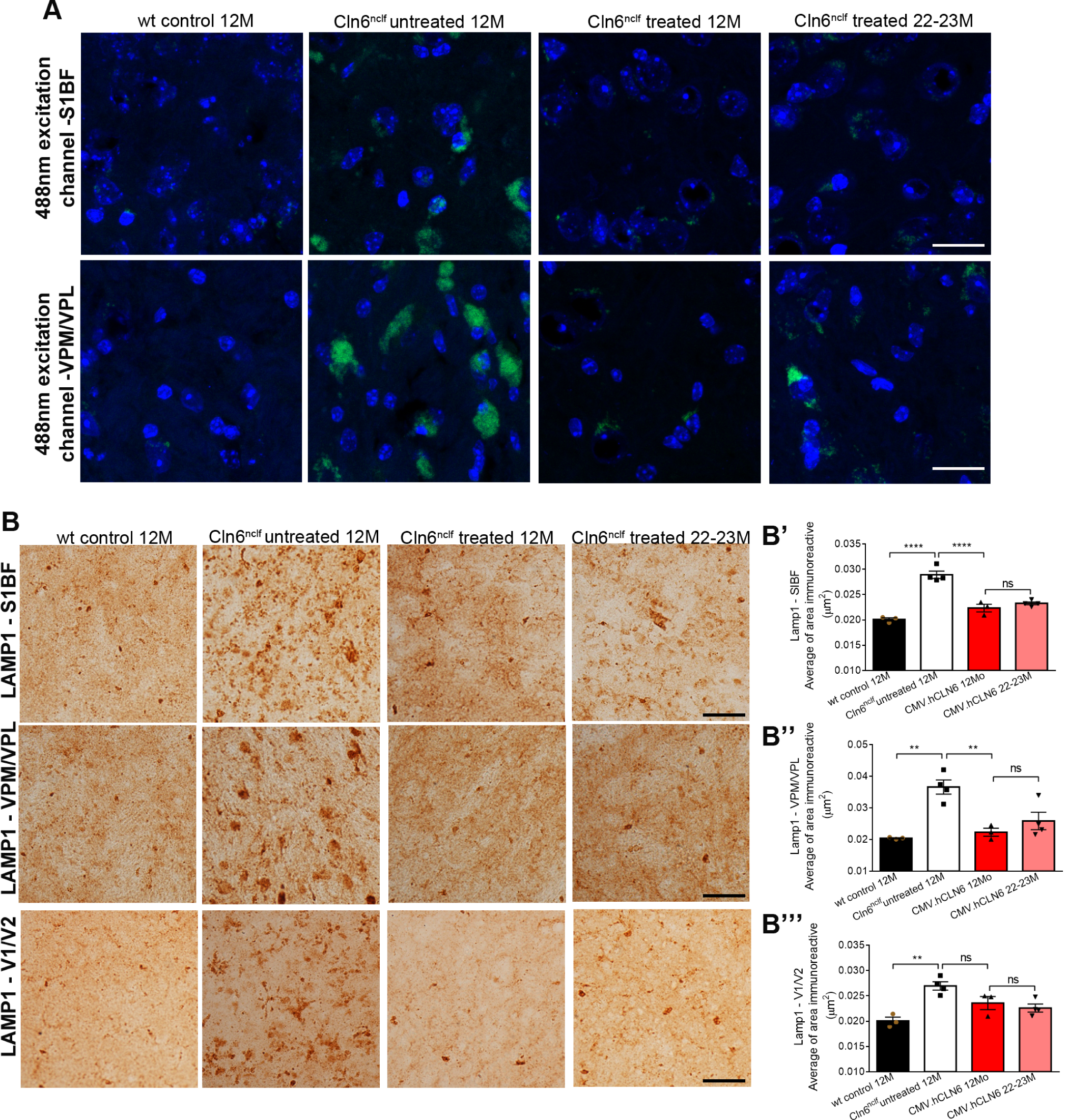
Brain-directed gene therapy preserves neuronal cells in *Cln6*^*nclf*^ mice long term. (A) Representative images of coronal brain cross-sections from 12-month-old end-stage *Cln6*^*nclf*^ mice, 12-month-old wild type mice, and 12- and 22-23-month-old treated *Cln6*^*nclf*^ mice. (B, C) Higher magnification images from S1BF and VPM/VPL regions stained for NeuN. (B’) Measurement of the cortical thickness in the S1BF region (B”, C’) and neuronal counts in the S1BF and VPM/VPL regions showing a significantly higher number of neurons in brains of *Cln6*^*nclf*^ mice at 12 months following neonatal ICV injections of vector. No significant difference was detected in the number of neurons in the S1BF and VPM/VPL region between brain sections of treated mutant mice aged 12 and 22-23 months. Scale bars: (A) 1 mm, (B, C) 50 µm. Wild type controls (12 months), n = 4; untreated *Cln6*^*nclf*^ (12 months), n = 4; treated *Cln6*^*nclf*^ (12 months), n = 3; treated *Cln6*^*nclf*^ mice (22-23 months), n = 4. Data are presented as means ± SEM and analysed by one-way ANOVA and Sidak’s multiple comparisons test (*p < 0.05, **p < 0.01).

### Gene therapy attenuates the microglia- and astrocyte– mediated immune response in *Cln6*^*nclf*^ mice

Finally, we assessed the effect of gene therapy on the inflammatory response by staining brain sections for CD68 and GFAP to label microglia and fibrillary astrocytes. We observed an increase in CD68-positive immunostaining in the cortex and the thalamus of untreated end-stage *Cln6*^*nclf*^ mouse brains compared with age-matched wild type controls and treated brains at 12 and 22-23 months of age (Figure 7A). In contrast with age-matched wild type controls where microglia are in a resting ramified state, activated and engorged CD68-positive microglia were present in S1BF, VPM/VPL and V1/V2 regions of untreated end-stage *Cln6*^*nclf*^ mice, as previously reported (9). Treated *Cln6*^*nclf*^ mice showed lower levels of CD68 staining in the S1BF, VPM/VPL and V1/V2 regions at 12 and 22-23 months (Figure 7B). Using thresholding image analysis, CD68 staining was quantified in all three brains regions. Significantly more immunoreactivity was present in untreated brain sections than in wild type controls at 12 months (p = 0.0025 in S1BF; p = 0.0131 in VPM/VPL; p = 0.0075 in V1/V2). The brain sections from treated *Cln6*^*nclf*^ mice had significantly less CD68 staining at 12 months than untreated end-stage controls (p = 0.0264 in S1BF; p = 0.0313 in VPM/VPL; p = 0.0217 in V1/V2). No significant differences were detected between the immunoreactivity in treated mice that were kept for 12 and 22-23 months post vector administration. The level of the CD68 immunostaining signal was comparable in the S1BF and the V1/V2 cortical regions of treated *Cln6*^*nclf*^ mice at 22-23 months and wild type mice at 12 months (Figure 7B’-B”‘).

**Fig. 7.**
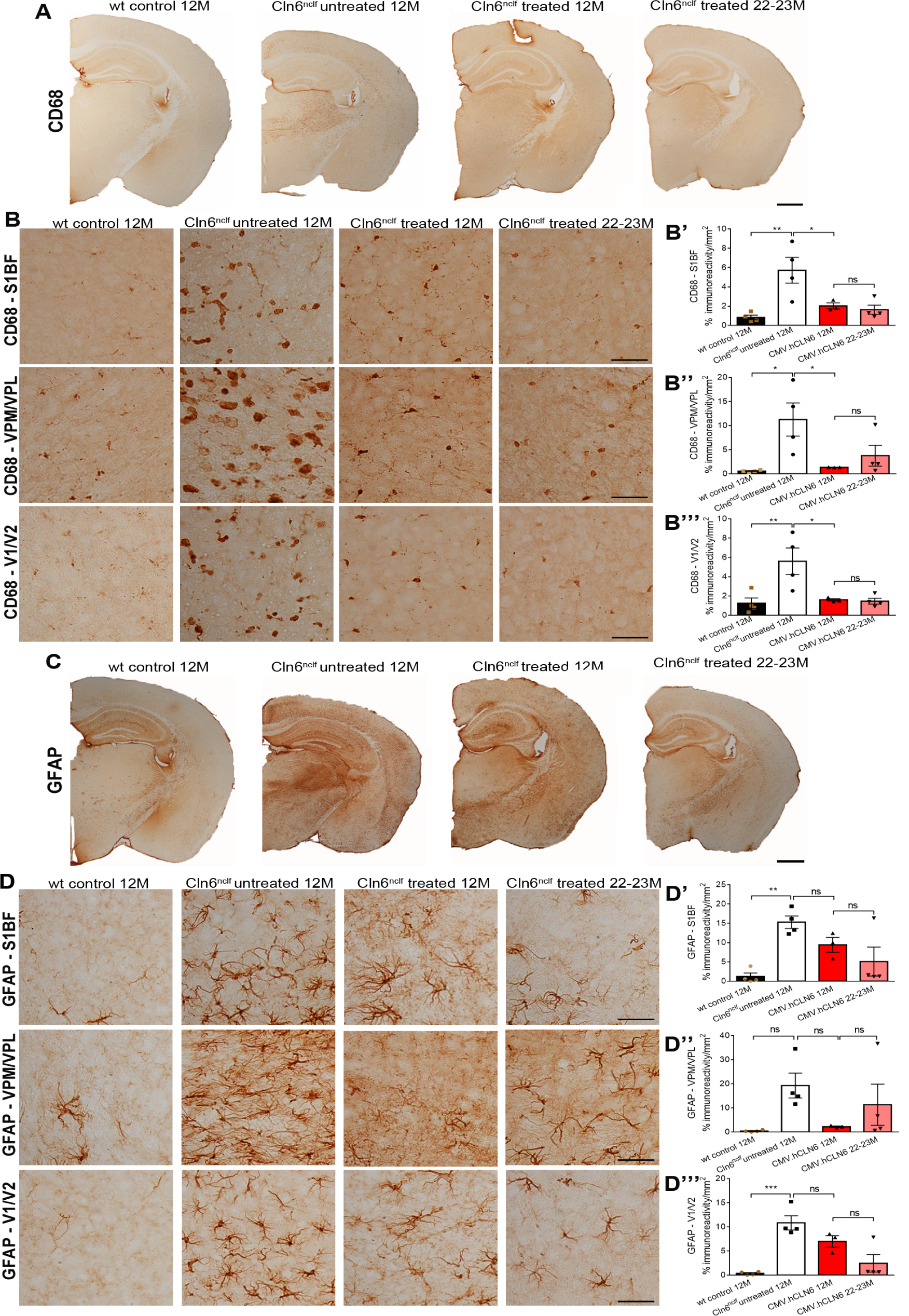
Brain-directed gene therapy preserves neuronal cells in *Cln6*^*nclf*^ mice long term. (A) Representative images of coronal brain cross-sections from 12-month-old end-stage *Cln6*^*nclf*^ mice, 12-month-old wild type mice, and 12- and 22-23-month-old treated *Cln6*^*nclf*^ mice. (B, C) Higher magnification images from S1BF and VPM/VPL regions stained for NeuN. (B’) Measurement of the cortical thickness in the S1BF region (B”, C’) and neuronal counts in the S1BF and VPM/VPL regions showing a significantly higher number of neurons in brains of *Cln6*^*nclf*^ mice at 12 months following neonatal ICV injections of vector. No significant difference was detected in the number of neurons in the S1BF and VPM/VPL region between brain sections of treated mutant mice aged 12 and 22-23 months. Scale bars: (A) 1 mm, (B, C) 50 µm. Wild type controls (12 months), n = 4; untreated *Cln6*^*nclf*^ (12 months), n = 4; treated *Cln6*^*nclf*^ (12 months), n = 3; treated *Cln6*^*nclf*^ mice (22-23 months), n = 4. Data are presented as means ± SEM and analysed by one-way ANOVA and Sidak’s multiple comparisons test (*p < 0.05, **p < 0.01).

GFAP immunostaining, followed by thresholding imaging analysis demonstrated widespread astrogliosis in untreated end-stage *Cln6*^*nclf*^ mouse brains compared with age-matched wild type controls. Greater GFAP-positive immunostaining was consistently observed in S1BF, VPM/VPL and V1/V2 regions of untreated *Cln6*-deficient mice (Figure 7C-D). Brain sections from treated *Cln6*^*nclf*^ mice at 12 and 22-23 months appeared to show a moderate reduction of GFAP immunostaining across all the brain regions compared with untreated end-stage mutant brain sections (Figure 7C-D). However, upon quantification, none of these differences reached statistical significance (Figure 7D’-D”’). These data indicate that gene therapy had beneficial effects on the brain pathology of treated mutant mice but the treatment did not prevent an inflammatory immune response entirely.

## Conclusions

Neurodegenerative LSDs such as CLN6 disease caused by mutations in genes that encode for membrane-bound proteins are more difficult targets for gene therapy than those conditions involving defective soluble enzymes. This is because the therapeutic effect from gene delivery is not further amplified through cross-correction of neighbouring cells. Recently, AAV-mediated pre-clinical gene therapy studies targeting the brain have shown efficacy in mouse models of Niemann-Pick type C1 disease (22, 23) and mucopolysaccharidosis type IIIC (MPSIIIC) (24), two neurodegenerative LSDs that are not cross-correctable. We took the approach of administering AAV9.CMV.hCLN6 via ICV injections to maximise transduction efficacy and prioritise therapeutic efficacy in the brain. We demonstrate that neonatal bilateral ICV injections of AAV9.CMV.hCLN6 with a total dose of 1×10^11^ and 5×10^11^ vector genomes significantly extended the survival of *Cln6*^*nclf*^ mice by over 90% to 23 months of age, effectively restoring normal lifespan. Furthermore, a range of behavioural tests assessing motor function of the treated mutant mice revealed either a preservation of motor skills and coordination or a complete normalisation of any deficits for almost two years following the treatment. These data demonstrate that the neonatal treatment does not only prolong the lifespan but also increases the quality of life of the treated mutant mice, an important conclusion for potential future translation into the clinic. Using landmarks of brain pathology previously established in *Cln6*^*nclf*^ mice by Morgan et al. (9), we measured a reduction in neuropathological markers of the disease and a significant decrease of neuronal loss in the cerebral cortex and the thalamus following AAV gene therapy, suggesting a sustained amelioration of neurodegeneration. Lysosomal storage material and microglial-mediated inflammatory response were reduced at 12 and 22-23 months. Although an overall trend towards amelioration of astrogliosis was observed in some regions, no statistically significant decrease of the astroglial-mediated inflammatory response was detected.

Our data demonstrated neonatal brain-directed AAV-mediated gene therapy successfully treats major features of CLN6 disease. However, our study also highlights some observations that require further investigation. Treated *Cln6*^*nclf*^ mice were more active in a short open field test as measured by an increased distance travelled and longer, more frequent visits to the centre of the test arena than age-matched wild type controls and untreated *Cln6*-deficient mice. It is possible that our treatment was not completely effective and uncovered a propensity for hyperactivity in *Cln6*-deficient mice, a phenotype that has thus far remained undetected due to a loss of mobility in the untreated animals. This may reflect increased anxiety or other neurological deficits. Assessing the open field phenotype of untreated mutants at earlier time points might have revealed disease related changes in activity behaviour of the mice at pre-symptomatic stages, which might clarify our observation. In the absence of a good antimouse Cln6 antibody, it was difficult to verify the endogenous expression pattern and level of *Cln6* in the neural cell types of the mouse brain. It is possible that our treatment failed to restore sufficient levels or resulted in inappropriate expression of *CLN6* for instance in astrocytes. Moreover, ICV administration of AAV9 leads to a widespread transduction of brain regions near the ventricles and the surface of the brain, but deep brain structures (e.g. the brain stem) were more sparsely transduced. It cannot be excluded that the increased activity could also be an adverse effect of a local *CLN6* overexpression. However, as we did not detect any other adverse effects of vector administration in treated mutant brains it seems unlikely that the hyperactivity may be caused by the transgene overexpression. Whilst it remains unclear why treated *Cln6*^*nclf*^ mice showed an increased activity, these findings emphasize the importance of thorough pre-clinical animal studies prior to clinical translation.

In our study the therapeutic vector was administered to the brains of *Cln6*^*nclf*^ mice at an early, pre-symptomatic stage of the disease. While neonatal vector administration is relevant to families with affected children that have undergone genetic testing and received diagnosis before disease onset, the more common clinical scenario is that a therapeutic vector would be administered upon symptomatic diagnosis. Moreover, we administered a volume of 10µl of AAV at 5×10^11^ and 1×10^11^ vg, a total vector dose that is difficult to meet in a clinical setting. Consequently, it will be important to assess the therapeutic effects of AAV gene therapy for CLN6 disease in older, symptomatic animals, and to assess the feasibility of widespread encephalic vector administration in larger animals. Some insight into the potential future translation of our approach for NCLs can be gained from a recent preclinical gene therapy study using a sheep model of CLN5 disease with a deficiency in the soluble Cln5 protein. Bilateral ICV injections with self-complementary AAV9.CB.Cln5 in symptomatic Cln5 sheep halted disease progression. These animals lived longer than untreated sheep, yet not as long as sheep treated pre-symptomatically, indicating that gene therapy administered via ICV injections after the onset of disease symptoms can still provide meaningful benefit (25).

Importantly, our study provides the first pre-clinical data set establishing that brain-directed gene therapy rescues the behavioural and neuropathological deficits in *Cln6*-deficient mice. In the past, AAV-mediated gene therapies have been reported to partially correct features of the disease phenotype in a model of CLN3 disease, which similar to CLN6 disease is caused by a defective transmembrane protein. Sondhi and colleagues showed a partial restoration of the neuropathological phenotype following multiple intracranial injections of AAVrh.10.CAG.hCLN3 in neonatal Cln3^ex7-8^ mice (26). Bosch et al. demonstrated that in young adult *Cln3*^*ex7-8*^ mice intravenous administration of scAAV9 carrying *CLN3* under the control of a weak neuronal promoter corrected the neuropathology up to 5 months post vector administration (27). To date, no long-term data from this study have been published. The authors also claim to rescue an early, non-progressive rotarod phenotype. However, other studies assessing rotarod performance in *Cln3*^*ex7-8*^ mice (28) and a second *Cln3*-deficient mouse strain (29) have not identified this phenotype. Together, these studies highlight the difficulties associated with developing brain-directed gene therapies for NCLs using models with a mild neurodegenerative phenotype. Here we have shown that a brain-directed gene therapy is therapeutic in pre-symptomatic *Cln6*^*nclf*^ mice, an NCL model with an earlier onset and a more severe disease progression. Our results also support ICV administration of AAV9-based gene therapy vectors as a promising strategy for other forms of NCL caused by transmembrane protein deficiencies.

## ACKNOWLEDGEMENTS

This project was supported by the Batten Disease Family Association, charity No 1084908, the European Union’s Horizon 2020 research and innovation programme (Grant No 66691), RP Fighting Blindness (Grant GR576), and Medical Research Council (Grant No MR/R025134/1). RRA is partially supported by the NIHR Biomedical Research Centre at Moorfields Eye Hospital and the UCL Institute of Ophthalmology.

## Material and Methods

### Plasmid construct and recombinant AAV production

To produce single stranded AAV9.CMV.hCLN6, the hCLN6 cDNA was cloned into a pD10 vector containing the CMV promoter as reported (19). Briefly, recombinant AAV2/9 was produced through a triple-transient transfection method and purified by size separation on a Sephacryl S300 column followed by anion exchange chromatography using a POROS 50 HQ column as previously described (30). Viral genome titers were determined by quantitative real-time PCR using a probe-based assay aligning to the SV40 poly-adenylation signal. Amplicon-based standard series of known amounts were used for sample interpolation. Final titres were expressed as total viral genome (vg) per pup.

### Neonatal intracerebroventricular (ICV) injections

*Cln6*^*nclf*^ mice were kindly provided by Prof Thomas Braulke (UKE Hamburg) and C57BL/6J mice were purchased from Harlan Laboratories (UK). Animal experiments were performed in accordance with the UK Home Office licence (PPL 70/8120) and according to ARRIVE guidelines and recommendations. At the age of post-gestational day (P) P0-P1 (under 24 hours) mouse pups received bilateral ICV injections of 5µl of vector into each lateral ventricle by using a 33-gauge needle (Hamilton, USA) as described previously (31).Following the injections, the neonates were subsequently returned to the parent cage

### Animal behavioural analysis

A range of behavioural tests was conducted to assess the performance of wild type, untreated *Cln6*^*nclf*^ and treated *Cln6*^*nclf*^ mice. When automated video analysis was not possible, researchers performing quantifications were masked to each experimental cohort of mice.

#### Accelerating rotarod

2-month-old mice were trained on 3 consecutive days with at least 2 trials per day on an accelerating rotarod (4 rpm to 45 rpm over 5 mins, Harvard apparatus, UK). A training trial was considered successful when the mice stayed on the rod for at least 45 sec. The resting period between trials was at least 15 mins. After successful completion of the training, the mice were tested on one day per month in two trials with a resting period in between the trials. The latency to fall was measured and the trial with the better performance was used for quantification.

#### 1-minute foot-fault test

Mice were placed on a stainless-steel grid (35×40 cm) with a mesh size of 1.3×1.3 cm, elevated 30cm above the bench surface. The animals were let to explore the grid for 1 minute. The number of steps and the number of times a limb fell through the grid (a foot-fault mistake) was counted to calculate the percentage of incorrect steps as described previously (32).

#### Vertical pole climbing

Pole climbing was assessed using a 28 cm long, wooden vertical pole with a diameter of 1.4 cm. The mice were placed at the top of the pole facing downwards and were given a 1-minute window to descend the pole. At 2 months, the mice were trained on 3 consecutive days with at least 2 trials per day. A training trial was considered successful when the mice descended the pole in a coordinated manner. After successful completion of the training, every month the mice performed the test twice on one day with a resting period in between both trials. The performance was scored as follows: 1 = pass, mouse descended down the pole in a coordinated manner, 0 = fail, mouse slipped or fell down the pole. The scores were averaged for both trails per mouse and per time point.

#### 30-seconds tail suspension test

The mice were suspended by their tails approximately 20 cm from the bench and were recorded for 30 seconds with a camera facing the abdomen of the animals as reported (33). The clasping phenotype was scored as follows: 3 = both hind limbs extended out from the body, 2 = one hind limb is partially retracted for less than 50% of the time, 1 = both hind limbs partially retracted from more than 50% of the time, 0 = both hind limbs clearly retracted and touching the abdomen.

#### 5-minute open field test

To assess activity and mobility, the mice were placed in an open field arena with white walls (48×48 cm). The mice were left to explore the arena for 1 minute and subsequently the mice were recorded for 4 minutes. The automated ANY-maze software (Ireland) was used to analyse the mouse behaviour and calculate various parameters including speed (mean, average), distance, time mobile, time immobile and time spent in the centre of the arena. The arena was thoroughly cleaned with 70% ethanol in between each test.

### Humane end point of study

The humane endpoint for this study was determined using a neurological welfare scoring system that assessed animal grooming, food and water intake, respiratory function, natural behaviour, provoked behaviour and neurological signs and/or a loss of 10-15% of bodyweight. Mice were not maintained for longer than 2 years of age.

### Tissue collection and brain section histology

Terminal exsanguinations were performed by trans-cardiac perfusion with phosphate-buffered saline (PBS). Brains were carefully extracted and fixed in 4% paraformaldehyde (PFA) for 48h at 4°C washed with PBS and stored in 30% sucrose in PBS until cryo-protected. Brains were cryo-sectioned at a thickness of 40µm along the rostro-caudal axis at −20°C using a cryostat microtome. Immunohistochemical staining was performed to analyse transgene expression, neuropathological marker proteins and cell type specific marker proteins. Endogenous peroxidase activity was depleted by incubating sections in 1% H^2^O^2^ tris-buffered saline (TBS) for 30 mins followed by three washes in TBS, after which endogenous non-specific protein binding was blocked by incubation in 15% normal serum (Sigma) in TBS-T (TBS with 0.3% Triton X-100) for 30 mins. The sections were incubated overnight in primary antibody for hCLN6 (1:500, non-commercial antiserum against CLN6 as reported (4)), GFAP (1:2000, MAB3402, Millipore, Massachusetts, USA), CD68 (1:2000, MCA1957, AbD Serotech, Hemel Hempstead, UK) or LAMP-1 (1:500, ab25245, Abcam) diluted in 10% normal serum in TBS-T. The next day sections were washed three times in TBS and incubated in 10% normal serum in TBS-T with biotinylated secondary antibodies anti-rabbit, anti-rat or anti-mouse IgG (1:1,000, Vector Laboratories Inc., Burlingame, CA, USA) for 2 hours. Staining was visualised using Vectastain avidin-biotin solution (ABC, Vector Laboratories) and DAB (Sigma). The sections were subsequently mounted, dehydrated, cleared in histoclear (National Diagnostics, Hessle, UK) for 30 mins and finally coverslipped with DPX (VWR, East Grimstead, UK). A live video camera (Nikon, DS-Fil, Melville, New York, USA) mounted onto a Nikon Eclipse E600 microscope was used to take representative images. To assess cell tropism and lysosomal storage material following the administration of AAV9.CMV.hCLN6, sections were stained as described above, with primary antibodies for CLN6 (1:500), NeuN (1:500, MAB377, Millipore), GFAP (1:2000, MAB3402, Millipore, Massachusetts, USA) or CD68 (1:100, MCA1957, AbD Serotech, Hemel Hempstead, UK). Secondary antibody was goat anti-rabbit Alexa488, goat anti-mouse Alexa568 or goat anti-rat Alexa568 (1:200, Life Technologies, Paisley, UK). Following three TBS washes, sections were counterstained with DAPI for nuclear visualisation, mounted onto glass slides and coverslipped. Sections were visualised with a laser scanning confocal microscope (Zeiss LSM 710, Carl Zeiss AG, Cambridge, UK).

### Quantitative analysis of immunohistochemical staining

To measure the levels of immunohistochemical staining, quantitative thresholding image analysis was used as previously described (34). For the quantification of GFAP- and CD68-postive immunostaining levels, 10 non-overlapping images were captured of each region of interest using a live video camera (Nikon, DS-Fil) mounted onto a Nikon Eclipse E600 microscope at x40 magnification with constant light intensity. For the quantification of the area occupied by LAMP1-positive accumulations, 6 to 10 non-overlapping images were captured using the same equipment and conditions as described above. Images were analysed using Image-Pro Premier (Media Cybernetics, Cambridge, UK). Immunoreactivity was measured using a constant threshold that was applied to all images for each respective antigen. Data are presented as the mean percentage area of CD68- and GFAP-positive immunoreactivity (± SD) and as the mean area of LAMP1 immunoreactive accumulations (± SD) for each region investigated. LAMP1-positive accumulations with an area below 0.01 µm^2^were discarded. To quantify the number of transduced cells also positive for NeuN, GFAP and CD68 in cortical areas manual counts were performed using the Image J software.

### Stereology

Measurement of cortical thickness and quantification of neuronal cell numbers were carried out on 40µm NeuN-stained mouse brain sections using StereoInvestigator software (MBF Bioscience, Williston, VT, USA). Images were acquired on a Nikon Optiphot microscope (Nikon) attached to a Q-Imaging Model 01-MBF-2000R-CLR-12 camera (MBF Bioscience). Cortical thickness of the somatosensory barrelfield cortex (S1BF) region was obtained by calculating the average of 10 perpendicular line measurements from the pial surface to the corpus callosum white matter on every sixth sections within the S1BF. Data are presented as the average of the cortical thickness (±SEM) per animal. Neuronal cell numbers within the S1BF and VPM/VPL were estimated using the optical fractionator probe. A border was traced around the region of interest, a grid was superimposed and neurons were counted within a series of 50µm × 50µm dissector frames, which were arranged according to the sampling grid size. Every sixth section containing the region of interest was analysed and NeuN-positive cells were counted at 100× objective. A coefficient of error (CE) between 0.05 and 0.1 was obtained for all counts indicating sufficient sampling efficiency. Data are presented as the mean of the estimated number of neurons (±SEM) per region and animal.

### Real-time qRT-PCR

Total RNA was extracted from distinct mouse brain regions using the RNeasy Mini Kit (QIAGEN). The QuantiTect Reverse Transcription Kit (QIAGEN) was utilized to generate cDNA. qRT-PCRs were performed in 96-well plates in a PCR thermal cycler (Applied Biosciences 7900HT) using a 2x FastStart TagMan Probe Mastermix assay (Roche) with a probe concentration of 100 nm and a primer concentration of 200 nm with the following primers: forward b-actin (5’-AAGGCCAACCGTGAAAAGAT-3’), reverse b-actin (5’-GTGGTACGACCAGAGGCATAC-3’), forward mouse Cln6 (5’-CGGGGACTACTTTCACATGAC-3’), reverse mouse Cln6 (5’-GGGGGACCGCTCAATAAG-3’), forward human CLN6 (5’-ATCACGCCCTTTCTCTTGC-3’), reverse human CLN6 (5’-TTGACAGAGTCACCCACCAG-3’), Mouse Cln6 primers were used for wild type mouse brain samples, human CLN6 primers were used for untreated and treated mouse brain samples and b-actin primers were used for all samples as loading control. Standard curves with human and mouse CLN6 cDNA ranged from 50 pg/µl to 0.05 pg/µl. Duplicate reactions were carried out for each sample. The Ct values were calculated using the computer program SDS 2.2.2 (Applied Biosciences). The obtained Ct values were averaged per duplicate reaction, and the number of specific cDNA molecules per nanogram total mRNA was calculated by standard curve intrapolation.

### Statistical analysis

Data are presented as means ±SEM or ± SD with the indicated n-numbers representing independent animals. GraphPad Prism 5 software (USA) was used to calculate significance and the appropriate tests and p values are provided in the individual figure legends.

---

**Fig. S1.**
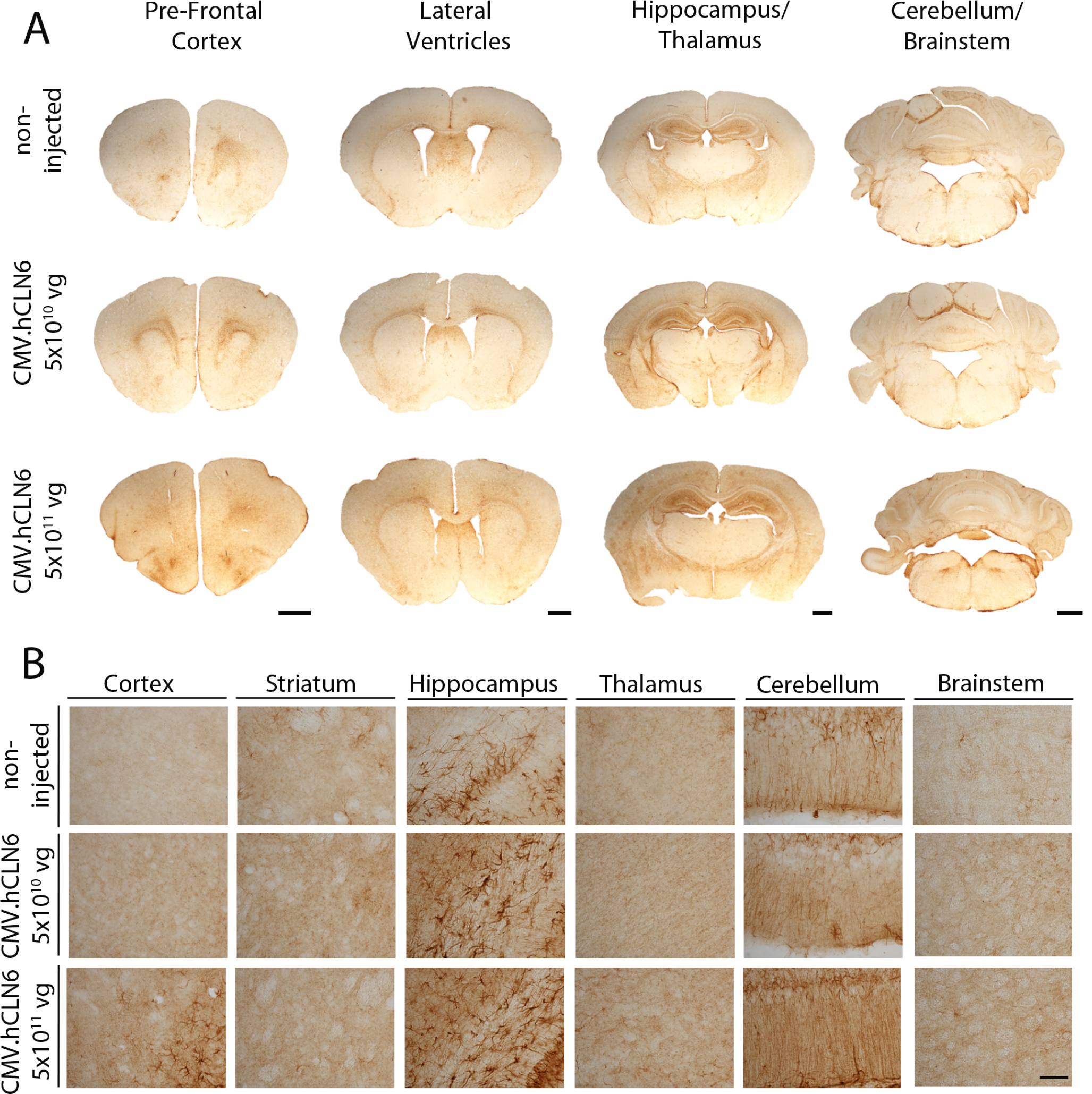
Overexpression of *CLN6* results in mild but localised astrocyte-mediated immune response in wild type mice. (A) Anti-GFAP immunostaining did not show an increase in astrogliosis in whole brain sections of wild type mice following neonatal ICV injections with AAV9.CMV.hCLN6 at 5×10^11^ vg. In comparison with non-injected age-matched wild type controls, a mild up-regulation of GFAP was detected in the pre-frontal cortex when AAV9.CMV.hCLN6 was administered at 5×10^11^ vg. (B) Higher magnification images confirmed a mild and confined increase in the astroglial-mediated immune response in the cortex of wild type brains following the administration of 5×10^11^ vg vector. Lower titre administration did not lead to increased astrogliosis across all brain regions assessed. Scale bars (A): 1mm, (B): 100 µm.

**Fig. S2.**
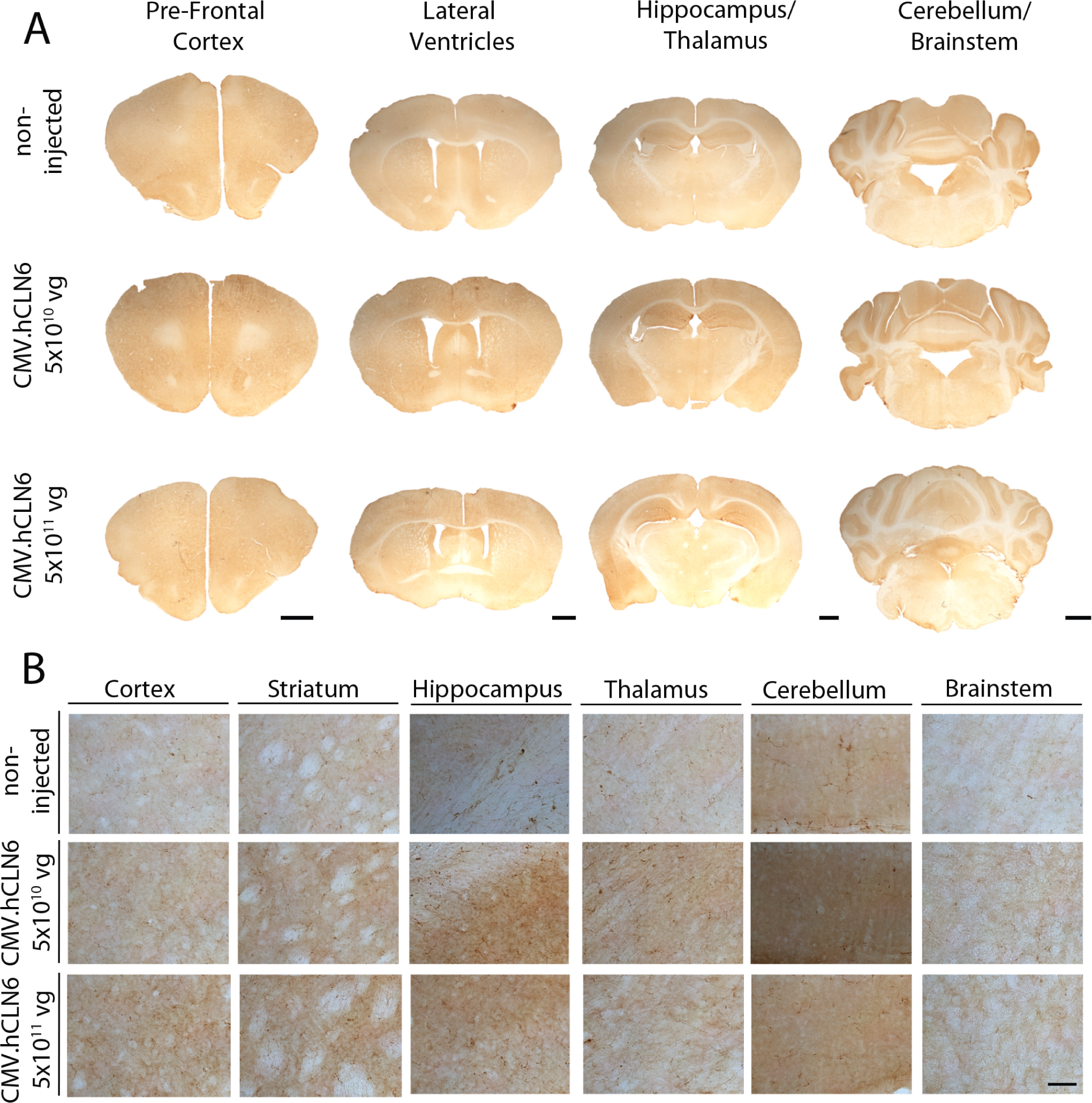
Overexpression of *CLN6* did not induce microglia-mediated immune response in wild type mice. (A-B) Anti-CD68 immunostaining did not reveal an increase in microglia activation across different brain regions following ICV administration of AAV9.CMV.hCLN6 at 5×10^10^ vg or 5×10^11^ vg in wild type mice compared with non-injected age-matched wild type controls. Scale bars (A): 1mm, (B): 100 µm.

**Fig. S3.**
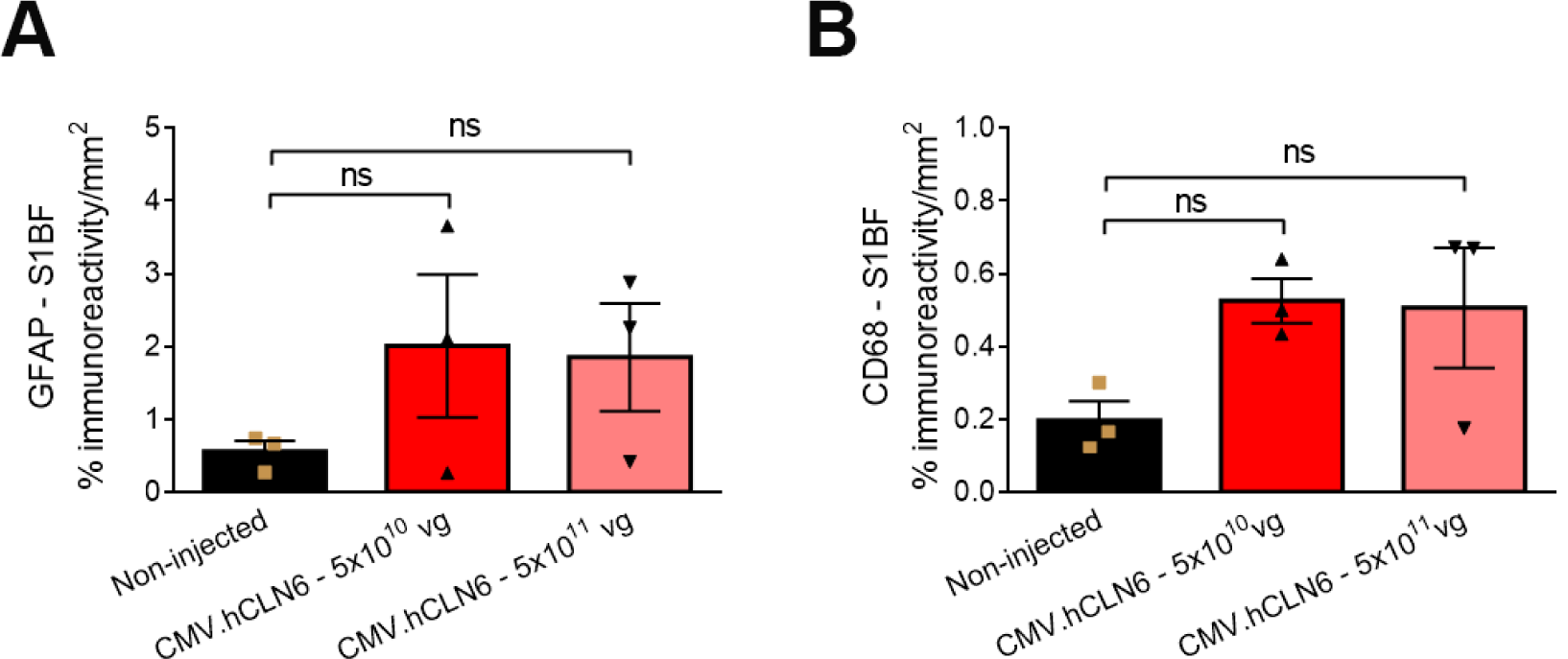
Quantification of the GFAP and CD68 immunoreactivity in the S1BF region of treated wild type mice. Wild type mice that received injections with AAV9.CMV.hCLN6 at 5×10^10^ vg and 5×10^11^ vg did not show significantly increased GFAP or CD68 immunoreactivity (B) in the S1BF region compared with untreated wild type controls. Non-injected controls, n = 3; 5×10^10^ vg, n = 3; 5×10^11^ vg, n = 3. Data are represented as means ± SEM, compared by one-way ANOVA.

**Fig. S4.**
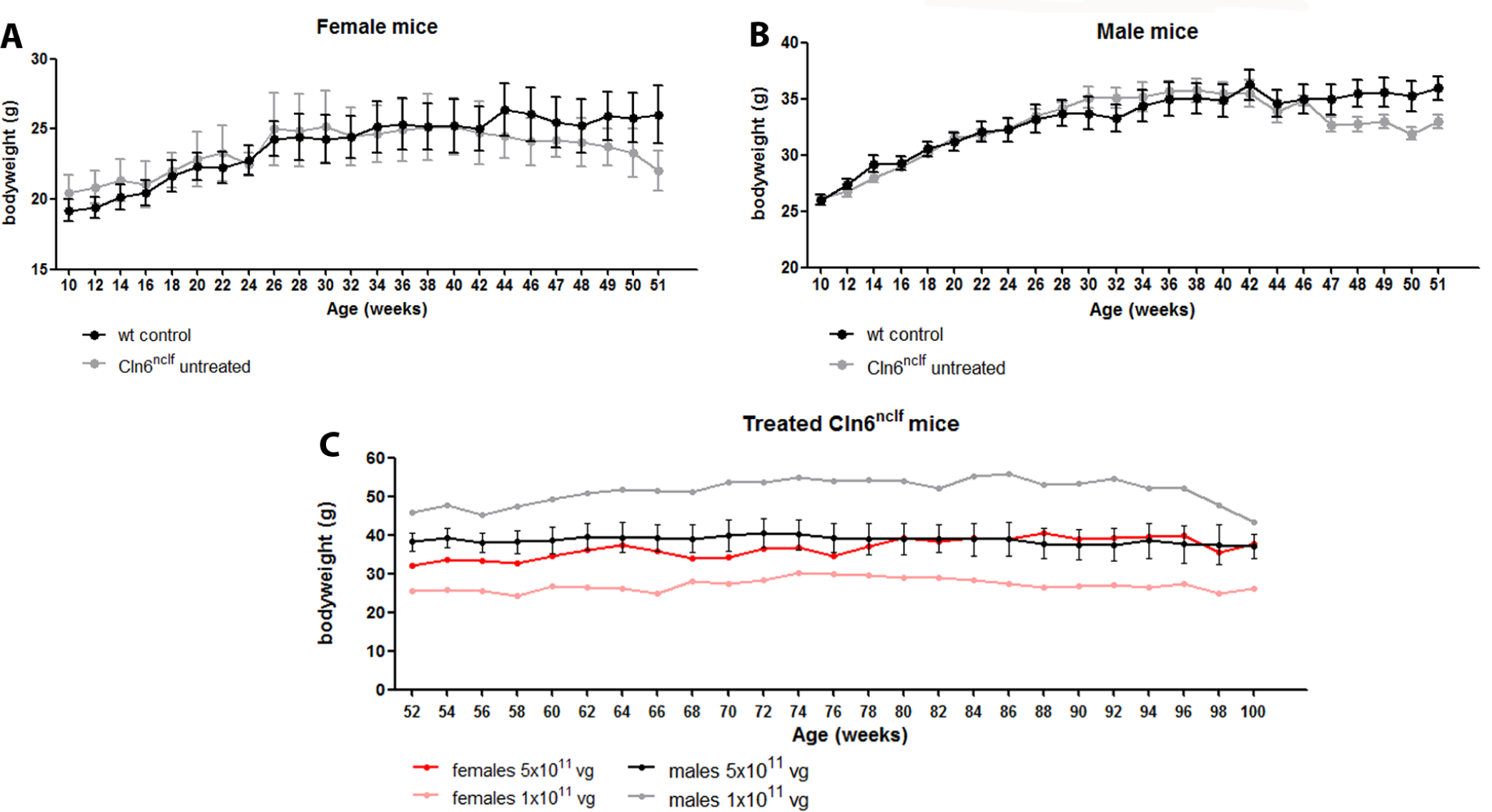
Bodyweights of wild type, untreated and treated *Cln6*^*nclf*^ mice over time. Bodyweights of (A) female wild type and *Cln6*^*nclf*^ mice, (B) male wild type and*Cln6*^*nclf*^ mice and (C) female and male AAV9.CMV.hCLN6-treated *Cln6*^*nclf*^ mice over time. Female wild type, n = 4; female *Cln6*^*nclf*^, n = 6; male wild type, n = 4; male *Cln6*^*nclf*^, n = 7; female *Cln6*^*nclf*^ treated with 5×10^11^ vg AAV9.CMV.hCLN6 vector, n = 1; female *Cln6*^*nclf*^ treated with 1×10^11^ vg vector, n = 1; male *Cln6*^*nclf*^ treated with 5×10^11^ vg vector, n = 5; male *Cln6*^*nclf*^ treated with 1×10^11^ vg vector, n = 1. Data represented as means ± SEM.

**Fig. S5.**
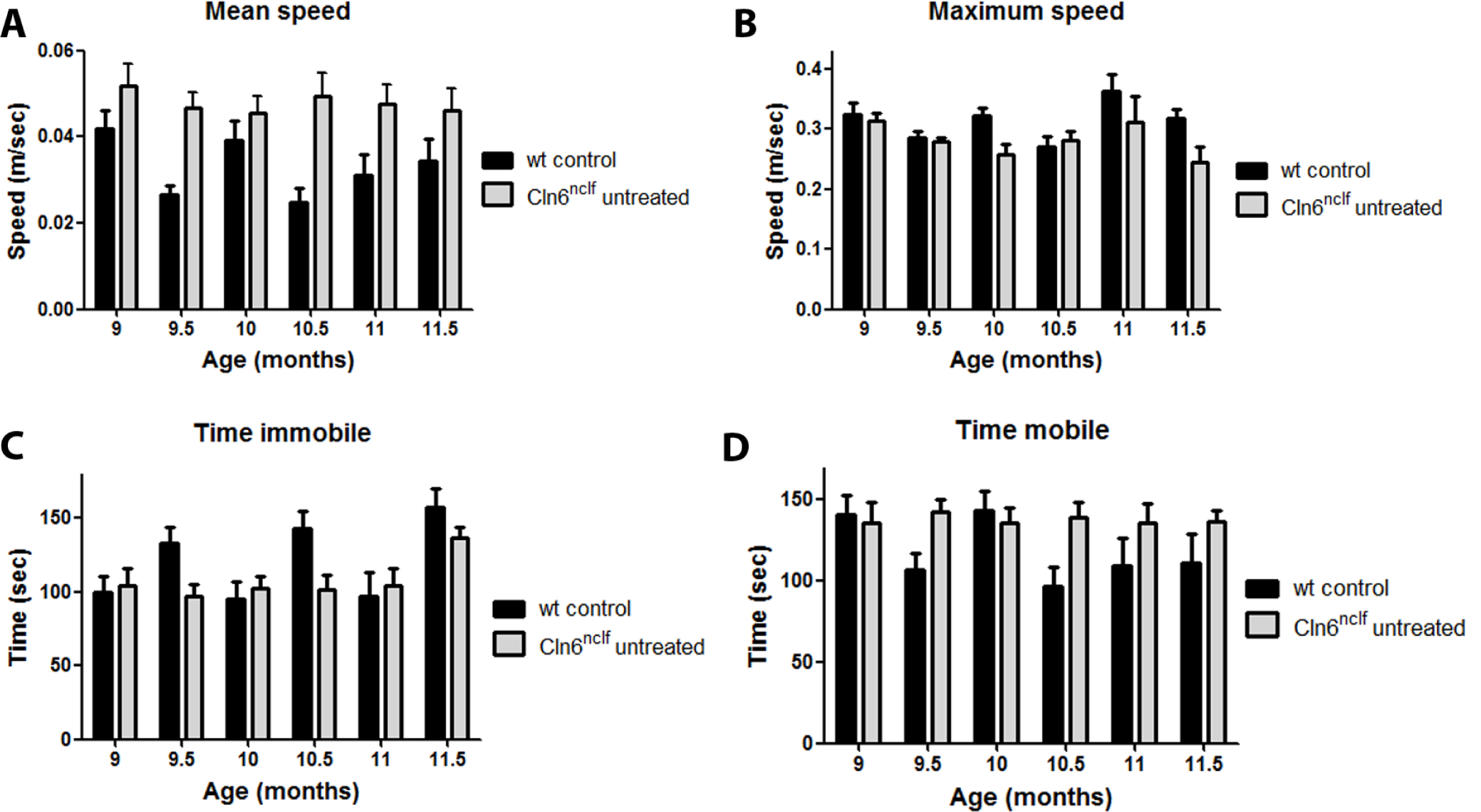
5-minute open field analysis of wild type and *Cln6*^*nclf*^ mice over time. Mean speed, (B) maximum speed, (C) time immobile and (D) time mobile were measured in a short 5-minute open field activity and mobility test. No significant differences were detected between *Cln6*^*nclf*^ and wild type mice over time. Wild type controls, n = 9-10; *Cln6*^*nclf*^ mice, n =11. Data are represented as means ± SEM, compared by two-way ANOVA.

**Fig. S6.**
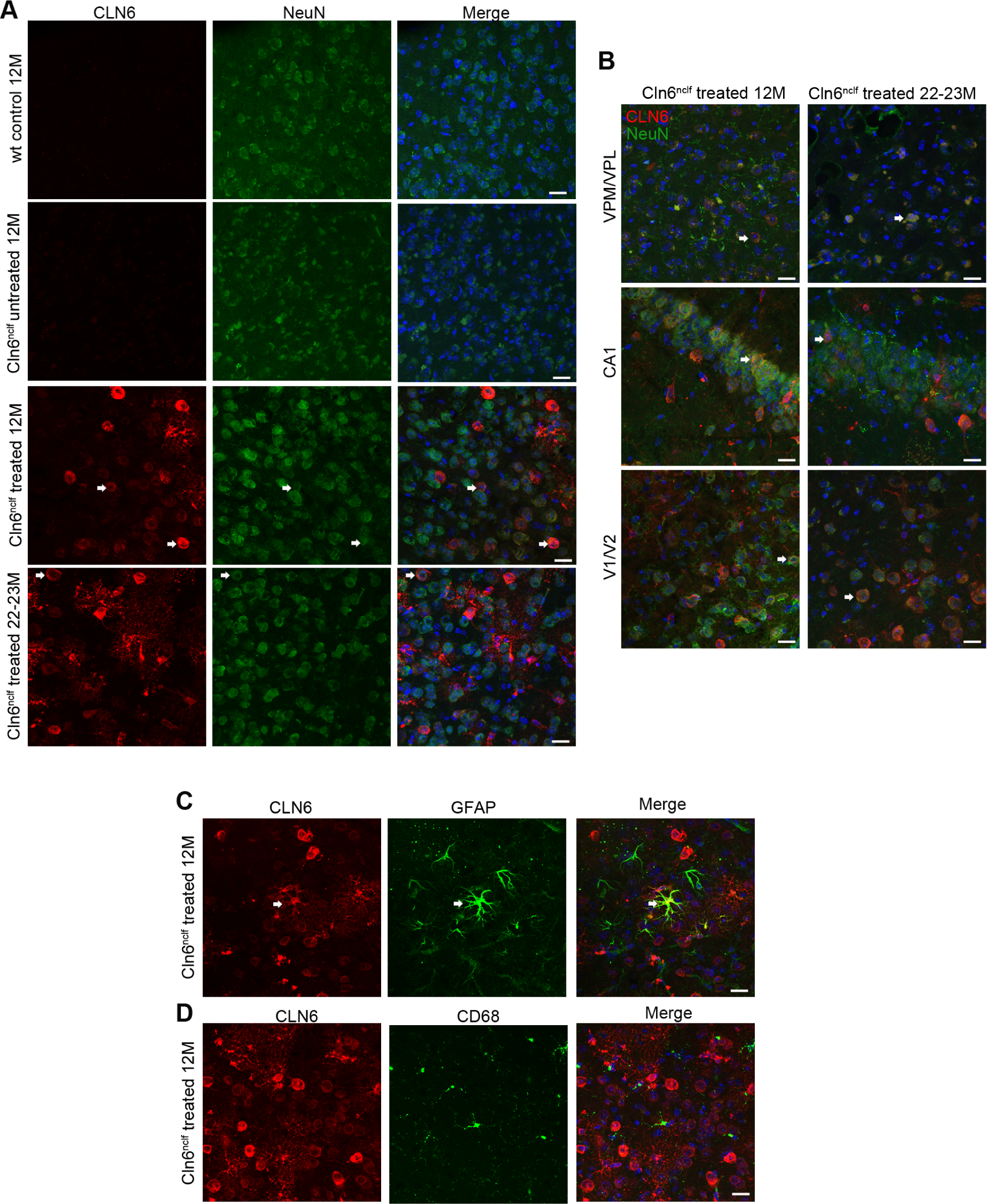
AAV9.CMV.hCLN6-treated mice show expression of *CLN6* in neurons and astrocytes. (A, B) Co-immunostaining with antibodies against CLN6 (red) and NeuN (green) confirmed that the *CLN6* transgene was expressed in S1BF, VPM/VPL, CA1 and V1/V2 region of treated *Cln6*^*nclf*^ mice at 12 and 22-23 months, co-localising with NeuN-positive cells (white arrows indicate examples). (C-D) Co-staining with antibodies against CLN6 (red), GFAP (green) and CD68 (green) demonstrating that the *CLN6* transgene was expressed in astrocytes (white arrow), only minimal expression was observed in microglia in treated brains. Scale bar: 20 µm

**Fig. S7.**
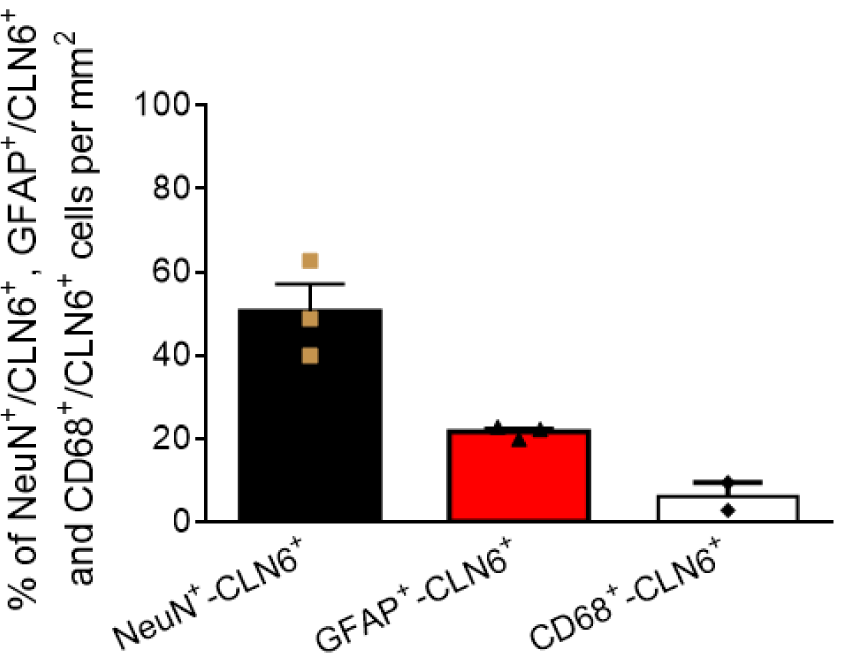
Transduction efficiency of NeuN-, GFAP- and CD68-positive cells following neonatal ICV administration of AAV9.CMV.hCLN6. To determine the percentage of transduced neurons, astrocytes and microglia, we counted the number of NeuN-, GFAP- and CD68-positive cells that were also positive for CLN6 staining in the cortical region of treated mutant brain sections using the software image J. We found that in 12 months old brain sections approximately 50 percent of NeuN-, 20 percent of GFAP- and less than 5 percent of CD68-positive cells expressed CLN6. NeuN^+^/CLN6^+^, n = 3; GFAP^+^/CLN6^+^, n = 3, CD68^+^/CLN6^+^, n = 2. Data are represented as means ± SEM.

**Fig. S8.**
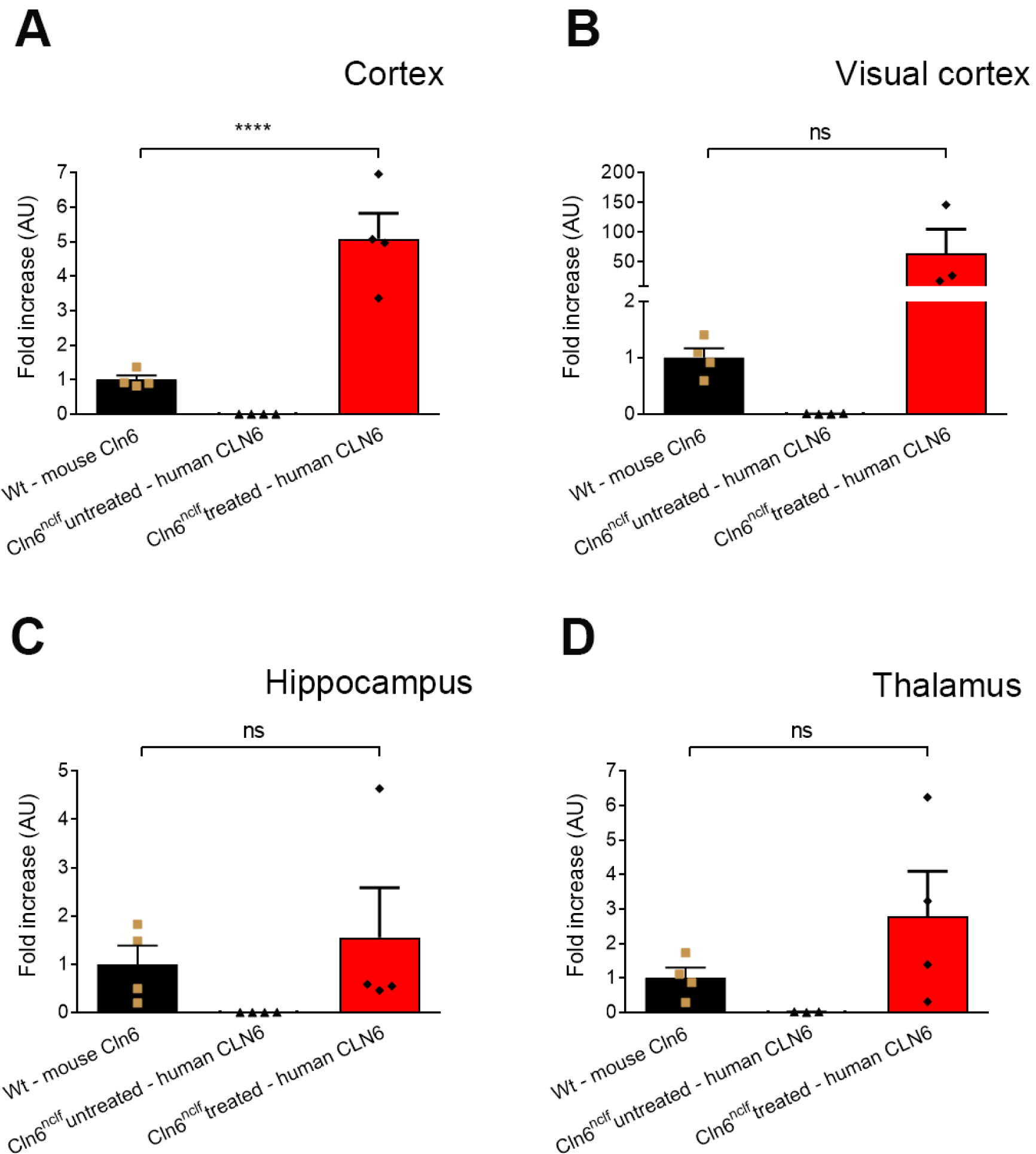
*CLN6* expression levels across different brain regions of wild type, untreated mutant and treated mutant mice at 12 months of age. A direct comparison of the absolute number of mRNA molecules by real-time qPCR showed that the CLN6 expression levels were significantly higher in the cortex of treated mutants than the endogenous Cln6 levels in the cortex of wild type mice (A). No significant difference was detected between wild type and treated mutants in the visual cortex (B), hippocampus (C) and thalamus (D). Data are represented as means ± SEM, compared by one-way ANOVA Sidak’s multiple comparisons test (****p < 0.0001).

